# A spiking neural-network model of goal-directed behaviour

**DOI:** 10.1101/867366

**Authors:** Ruggero Basanisi, Andrea Brovelli, Emilio Cartoni, Gianluca Baldassarre

## Abstract

In mammals, goal-directed and planning processes support flexible behaviour usable to face new situations or changed conditions that cannot be tackled through more efficient but rigid habitual behaviours. Within the Bayesian modelling approach of brain and behaviour, probabilistic models have been proposed to perform planning as a probabilistic inference. Recently, some models have started to face the important challenge met by this approach: grounding such processes on the computations implemented by brain spiking networks. Here we propose a model of goal-directed behaviour that has a probabilistic interpretation and is centred on a recurrent spiking neural network representing the world model. The model, building on previous proposals on spiking neurons and plasticity rules having a probabilistic interpretation, presents these novelties at the system level: (a) the world model is learnt in parallel with its use for planning, and an arbitration mechanism decides when to exploit the world-model knowledge with planning, or to explore, on the basis of an entropy-based confidence on the world model knowledge; (b) the world model is a hidden Markov model (HMM) able to simulate sequences of states and actions, thus planning selects actions through the same neural generative process used to predict states; (c) the world model learns the hidden causes of observations, and their temporal dependencies, through a biologically plausible unsupervised learning mechanism. The model is tested with a visuomotor learning task and validated by comparing its behaviour with the performance and reaction times of human participants solving the same task. The model represents a further step towards the construction of an autonomous architecture bridging goal-directed behaviour as probabilistic inference to brain-like computations.

**Author summary:** Goal-directed behaviour relies on brain processes supporting planning of actions based on the prediction of their consequences before performing them in the environment. An important computational modelling approach of these processes sees the brain as a probabilistic machine implementing goal-directed processes relying on probability distributions and operations on them. An important challenge for this approach is to explain how these distributions and operations might be grounded on the brain spiking neurons and learning processes. Here we propose a hypothesis of how this might happen by presenting a computational model of goal-directed processes based on artificial spiking neural networks. The model presents three main novelties. First, it can plan even while it is still learning the consequences of actions by deciding if planning or exploring the environment based on how confident it is on its predictions. Second, it is able to ‘think’ alternative possible actions, and their consequences, by relying on the low-level stochasticity of neurons. Third, it can learn to anticipate the consequences of actions in an autonomous fashion based on experience. Overall, the model represents a novel hypothesis on how goal-directed behaviour might rely on the stochastic spiking processes and plasticity mechanisms of the brain neurons.

## Introduction

In mammals, the acquisition and consolidation of instrumental behaviour involves two sets of processes, one underlying flexible *goal-directed behaviour*, used in particular to find solutions to new problems or face changing conditions, and the other one related to *habits*, forming stimulus-response behaviour used to efficiently but inflexibly face familiar conditions [1–3]. As also highlighted in the computational literature [4], goal-directed processes are *model based*; that is, they rely on an internal representation of the external world (*world model*) to internally simulate (*planning*) the consequences of actions, or action sequences, usable to achieve desired world states (*goals*) before executing them in the environment [4–7]. When the agent has a model of the relevant part of the world and has to accomplish a new goal, goal-directed behaviour allows it to solve the task on the basis of planning and the world model. This thanks to the fact that the world model represents the general dynamics of the world, in particular how it responds to the agent’s actions, and so it can be used to pursue any goal (in particular, goal independent). The simulated achievement of the new goal might be possibly marked by an internal reward [8]. To an external observer the agent appears to solve the new task ‘on the fly’ or ‘by ‘insight’. Instead, habitual behaviour is *model free*, in the sense that it relies on actions directly triggered by stimuli (*habits*) and does not require a world model anticipating their outcomes [4, 6, 9]. Habits are task dependent as they rely on stimulus-responses associations that can lead the agent to specific desired world states. Given a new desired state, the agent thus needs a repeated experience of such state to discover and learn by trial and error the new stimulus-response associations leading to it.

When a goal-directed system encounters a new task that involves an unknown part of the environment, or a part of the environment that changed, it first needs to learn a model of it (or to update the existing model) before using it for planning. In this respect, goal-directed behaviour involves two subsets of processes, which tend to characterise two successive phases when a new problem or a changed environmental condition are faced. The first subset of processes are directed to the *exploration* of the environment to form the internal model of it, while the second subset of processes are directed to the *exploitation* of the acquired knowledge to plan and execute actions successfully accomplishing the desired goal [10, 11]. Here we consider the early phases of the solution of a new task, involving either a new environment or a new goal, and hence we focus on goal-directed behaviour and its exploration/exploitation processes.

In brain, goal-directed behaviour relies on ventral/associative basal ganglia and frontal cortex supporting the anticipation of the world dynamics and action consequences; instead, habitual behaviour relies on motor basal ganglia and sensorimotor/premotor cortices able to acquire stimulus-response associations by reinforcement learning [9, 12–14]. The brain processes underlying goal-directed behaviour have been interpreted within different computational frameworks. A current influential view of brain, rooted in Helmholtz’ pioneering contributions on perception [15], considers it a *probabilistic* or *Bayesian machine* that copes with the uncertainties of the world by representing it in terms of probability distributions and probability inferences on them pivoting on the Bayes rule [16, 17]. This view of brain has been progressively extended to cover all aspects of cognition, from perception to action and decision making (e.g., [18, 19]).

Within this framework, it has been proposed that brain also implements goal-directed behaviour and planning through probabilistic representations and inferences, and this has been shown with specific models (e.g., [20–22]). These models rely on various probabilistic processes to represent the world, some of which are shown in Fig 1 through their corresponding graphical models.

**Fig 1.**
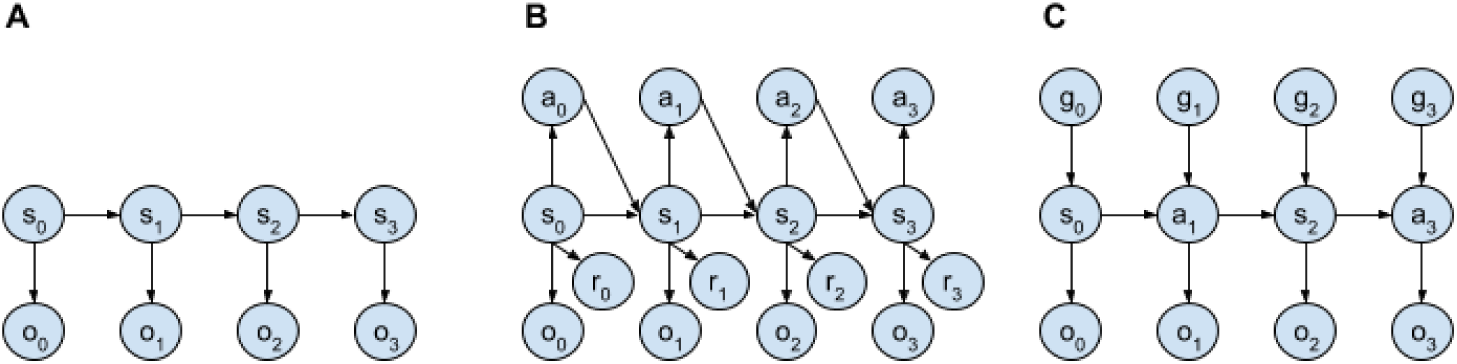
Graphical models of some probabilistic models usable to represent the dynamics of the world in planning systems. Nodes represent probability distributions and directional links represent dependencies between conditional probabilities. (a) Hidden Markov Models (HMM): these are formed by state nodes ‘s’ and observation nodes ‘o’. (b) Partially Observable Markov Decision Processes (POMDP): these are also formed by action nodes ‘a’ and reward nodes ‘r’ (different versions of these model are possible based on the chosen nodes and their dependencies). (c) The HMM considered here, where both states and actions are considered as ‘observed events’ by the planner.

*Hidden Markov Models* (HMM) are one important means used to represent the world dynamics [23, 24]. A HMM assumes that the agent cannot directly access the states of the world (they are ‘hidden’ to it) but only infer them on the basis of noisy information from sensors. The model thus internally represents the states of the world as probability distributions over possible *hidden causes* of observations, in particular with a different distribution for each time step. The probability distribution over states at each time step is assumed to depend only on the state of the previous time step (*Markov property*). The model also internally represents the probability distribution over the possible observations and assumes that it depends only on the current state. An agent can use a HMM representing the world dynamics to internally simulate possible sequences of states that the environment might traverse; the model is also *generative* in the sense that it can simulate the possible observations caused on the sensors by the internal activation of the internal representatios of the world states.

*Partially Observable Markov Decision Processes* (POMDP) again assume that the agent can access the states of the world only indirectly through noisy sensors (they are ‘partially observable’) but also consider the agent’s behaviour, in particular as probability distributions over actions at different times. The probability distributions over actions are conditioned on internal representions of states (thus forming probabilistic *policies*), and over perceived rewards. Rewards are considered as additional observations and as such are assumed to depend on other events such as the world states (different models can vary on their assumptions on rewards). POMDPs can be used to implement planning by conditioning probability distributions on a high reward (or a reached goal state), and then by inferring the probability distributions of the state-action sequences causing them with a high likelihood (*planning as inference* [20–22]).

Probabilistic models have the strength of capturing the uncertain nature of the world and the possible probabilistic representations and inferences that the brain might employ to represent them. However, their use as models of the brain, and not only of behaviour, encounters an important challenge; namely, the fact that the probability distributions that these models commonly use directly involve abstract/high-level aspects of cognition and behaviour, such as the probability distribution over world-states, actions, and observations, and so this opens up the problem of explaining how such distributions and the inferences on them could rely on the firing of brain neurons [17, 25, 26].

One important possibility is that the needed probability distributions could rely on the probability distributions of spikes of neurons, sampled by the actual spikes, and that the connections between neural populations, undergoing experience-dependent plasticity, supports the conditional probabilities underlying the needed probabilistic inferences [21, 27–30]. An interesting approach to implement this idea, on which we build here, proposes mechanisms to ground important building blocks of probabilistic models on spiking neural networks similar to those of the brain, on some typical connectivity patterns of cortex, and on biologically plausible plasticity rules [24, 31–33]. This approach proposes how spiking networks and spike-timing dependent plasticity (STDP) could model the learning of hidden causes of observations [32]. In particular, the spikes of neurons in different trials can represent probability distributions over internal representations of stimuli and a typical connectivity pattern found in cortex (the *winner-take-all* pattern relying on lateral inhibition), and STDP, can lead to the emergence of circuits able to identify the hidden causes of observations. Moreover, STDP can also support the formation of probabilistic dependencies between such hidden causes, capturing their relations in time. This can instantiate an HMM usable to internally represent the perception of *sequences* of events [24, 32, 33].

Recently, mechanisms as these have been used in recurrent spiking neural network models to implement planning [34–36]. These models are the state-of-the-art in the realisation of probabilistic models of planning grounded on biologically plausible spiking neural networks. In these models, a two-dimensional neural map of spiking neurons with lateral connectivity is used as a model of the world, with the world consisting in a scenario involving navigation or robot motion tasks. The lateral connectivity network implementing the model of the world, pre-trained with supervised learning, encodes information about which locations in space are linked to which other locations. The unconditioned spikes of the world model sample the prior probability of the state sequences followed by the agent if it explores the environment randomly, and of the rewards associated to the sequence (e.g., a reward of 1 when the target state is achieved). A second neural layer of spiking neurons that encodes ‘context’, intended as the task target state (e.g., in a navigation task), has all-to-all connections to the world model and can condition the probability distribution expressed by it. The neural solution to the inference problem relies on the update of the connections linking the context to the world model so that the distance (Kullback-Leibler divergence) between the prior probability distribution of the sequences converges to the desired posterior probability maximising the reward. The actions needed to follow the state sequences sampled from the posterior distribution are inferred, after the sampling, by inverse kinematics, either offline [35] or using a dedicated layer [36].

Here we propose a neural spiking architecture for probabilistic planning that builds on previous contributions, overcomes some of their limitations, and introduces some novelties relevant for modelling biologically plausible planning. At the low level, the model relies on the neural mechanisms and the unsupervised learning plasticity rule proposed in [24] (plus a reinforcement learning rule) to implement a HMM with a spiking recurrent neural network. A recurrent spiking neural network to implement the world model and planning was also used in [35, 36]. With respect to these previous models, our architecture presents a number of structure and functioning novelties at the system level.

A first novelty of the architecture is that the learning of the world model is intermixed with its use in planning, as it is required by the fact that goal-directed behaviour often tackles tasks involving new or changed parts of the world. This implies the non-trivial challenge that planning must be performed on the basis of a partial model of the world. To face this problem, the model uses an *arbitration* component, inspired by the mechanisms proposed in [37, 38] for the arbitration between goal-directed and habitual control, that decides between the *exploitation* of the world model knowledge and the *exploration* of the environment. To this purpose, the arbitration measures the uncertainty of the world model based on the entropy of its probability distributions. When this uncertainty is low, planning continues, otherwise the exploration component selects an action based on previous experiences or randomly.

A second novelty of the architecture is that the world model is a HMM that ‘observes’, learns, and predicts sequences formed not only by states but also actions. These actions are ‘observed’ by the world model when executed in the world after being selected by the planning process or the exploration component. This allows the model to predict states and actions through the same neural probabilistic generative mechanisms. Given the observation of the current world state, the world model produces a probability distribution over the state-action sequences that favour the event sequences actually observed in the world. Each sampling hence tends to select one of these sequences. The world model however also receives connections from a neural representation of the pursued goal. When the architecture experiences a successful achievement of the goal in the environment, those connections are updated so that the likelihood of selecting the sequence leading to the goal progressively increases.

A third novelty of the architecture is that the lateral connections within the world-model neural network, instantiating the probabilistic time dependencies between the hidden causes of the HMM, are learned in an *unsupervised* fashion, in particular on the basis of the STDP mechanism proposed in [24]. This is an advancement with respect to biological plausibility of the architecture, with respect to using supervised learning as done in previous spiking network models of goal-directed behaviour [35, 36], as the system is able to autonomously learn *internal* representations of the observed events and event-sequences without the need of an external ‘teacher’ suggesting them.

A last contribution of this work is the use of the architecture to reproduce and interpret the results of the experiment presented in [12] where human participants learn to solve a visuomotor learning task. This allowed the validation of the model, in particular to check if the learning processes of the world model lead to match human performance, and if the arbitration mechanism employing a variable time planning lead to reproduce the reaction times exhibited by human participants. This target experiment was also investigated by the model proposed in [38]. Although this model did not aim to bridge probabilistic modelling to neural mechanisms as here, it used an interesting mechanism of arbitration between goal-directed and habitual behaviour based on entropy as done here.

The rest of the paper is organised as follows. Section 1 describes the model architecture and functioning and the visuomotor learning task used to validate it. Section 2 presents and discusses the results of the model tests, in particular by comparing the model performance and reaction times with those of human participants of the visuomotor task, and by showing the mechanisms that might underlie such performance. Finally, Section 3 draws the conclusions.

## 1 Materials and methods

This section explains the model architecture and functioning and the visuomotor task used to test it [12]. Although the main objective of this work is to propose the novel spiking neural model of goal-directed behaviour, the section starts by illustrating the visuomotor task to use it as an example while illustrating the model.

### 1.1 Target experiment

In the task proposed in [12], the participants are supposed to discover the correct associations between three different stimuli and three, out of five, possible motor responses. During the experiment, three different colours are projected on a screen in a pseudo-randomised order, in particular through sixty triplets each involving each colour once in a random order. After each colour perception, the participants have to press one of the five buttons of a keyboard with their right hand. Once the action is performed, a feedback on the screen informs the participants if the association between the colour and the performed action was correct or wrong. Unbeknown to the participants, a fixed number of errors is used to dynamically consider the action performed at a certain time steps as correct for the particular colour: the correct action for S1 comes after one error (so at the second attempt), for S2 after three errors (fourth attempt), and S3 after four errors (fifth attempt). The activity of the participants is supposed to be organised in two phases: an initial exploratory phase where they the correct associations and a second exploitation phase where should repeat the found correct associations until the end of the task (Fig 2). The participants are thus not supposed to explore all the possible colour-action associations since their objective is to discover and exploit one correct association per colour.

**Fig 2.**
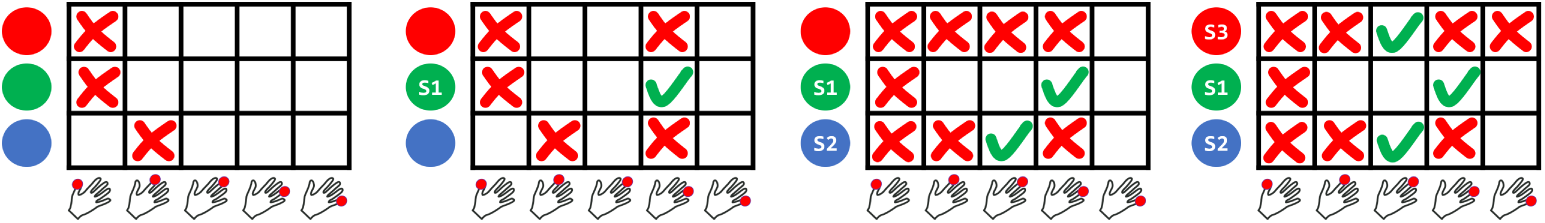
The visuomotor learning task used to validate the model. Three colour stimuli are presented to the participants in a pseudo-random order, in particular in triplets containing all three colours once in a random order. The action consists in pressing one out of five possible buttons with the right hand. The figure shows four triplets of an ideal participant that never repeats an error for a certain colour and does not forget a found correct action. The colour receiving the first action in the second triplet is marked as the first stimulus (S1), and such action is considered the correct one for it. The colour different from S1 receiving the first action in the fourth triplet is marked as the second stimulus (S2), and such action is considered the correct one for it. The colour different from S1 and S2 receiving the first action in the fifth triplet is marked as the second stimulus (S3), and such action is considered the correct one for it.

### 1.2 Goal-directed behaviour model: overview of the architecture and functioning

#### 1.2.1 Architecture

The model is composed of a spiking neural network for planning formed by four different layers, a spiking neural network for exploration formed by two neural layers, and an arbitration component. Fig 3 gives an overview of the structure and functioning of the architecture. The planning neural network instantiates a HMM and is formed by four layers of neurons now considered in more detail.

**Fig 3.**
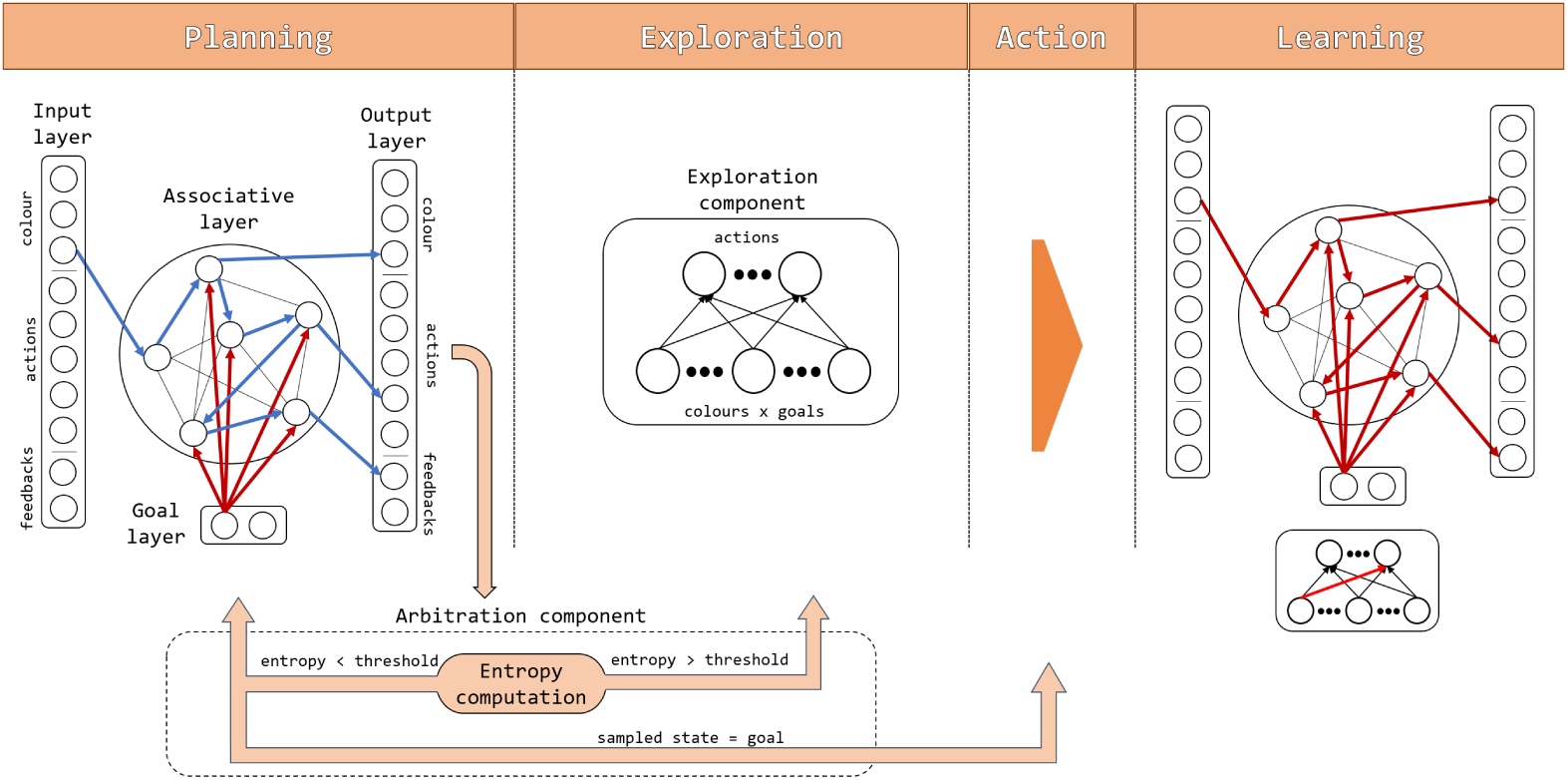
Components and information flow of the spiking neural-network architecture. Blue arrows represent information flows travelling stable connections between units while red arrows represent information flows travelling connections that are updated in the considered phase. The main loop of a single trial can be divided in four different phases: planning, (possibly) exploration, action execution, and learning (see text).

##### Input layer

The input layer contains ten neurons, three encoding the stimuli (colours), five encoding the actions (different possible finger presses), and two encoding the outcome (positive or negative feedback). The input layer sends all-to-all afferent connections to the neurons of the associative layer.

##### Goal layer

The goal layer can be composed of a variable number of neurons corresponding to the goals to achieve (two goals in the visuomotor task: ‘obtain a positive feedback’ and ‘obtain a negative feedback’). When the agent commits to a goal it activates its corresponding unit on the basis of internal mechanisms not simulated here. Goal neurons send all-to-all efferent projections to the associative neurons.

##### Associative layer

The associative layer, forming the core of the model, is composed of 400 neurons, all connected to each other but without self-connections. The associative layer receives the mentioned afferent connections from the input and goal layers, and sends all-to-all efferent connections to the neurons of the output layer.

##### Output layer

As the input layer, the output layer is composed of ten neurons each one representing one of the stimuli, actions, and outcomes of the task. The output layer receives the mentioned afferent connections from the associative layer.

Together the four layers instantiate a neural HMM implementing the system’s world model used for planning. In particular, the input and output layer together form the observation part of the HMM, and have an identical structure. Given the one directional nature of brain connections, we used the two layers to implement separately the two functions played by the observation part of the HMM, namely the input from the external environment and the possible generative reconstruction of such input based on internal causes. The associative layer implements the probability distribution over the hidden causes of the observations and the probabilistic temporal dependencies between them. The goal layer can condition the latter distributions to possibly increase, with learning, the probability of sampling simulated stimulus-action-outcome sequences that lead to the desired goal.

Alongside the planning components, the system was formed by the following additional components used for exploration and arbitration.

##### Exploration component

This component is also formed by two layers of spiking neurons: (a) the input layer encodes the combinations of stimuli and goals (3 × 2 neurons corresponding to 3 colours and 2 goals), and (b) the action layer composed of five neurons (the five possible finger presses).

##### Arbitration component

This component decides when to plan, to explore, or to act in the world. Currently the component has not a neural implementation. The decision is made on the basis of the knowledge of the world model, measured as the average entropy of its probability distribution during a planning cycle. When entropy is lower than a threshold, and a goal has not been found, planning continues, whereas if a goal has been found the corresponding action is performed in the environment. If entropy is above the threshold then the control is passed to the exploration component that selects the action to perform in the world.

### 1.2.2 Functioning

The functioning of the model is summarised in Algorithm 1. The model experiences multiple trials of the task: 60 (20 colour triplets) with the goal set to ‘achieve a positive feedback’ (this reflects the target experiments) and 60 (other 20 colour triplets) with the goal set to ‘achieve a negative feedback’ (we shall see these additional trials are used to produce a prediction of the model). A certain number of discrete time steps of the simulation (here 15) is assumed to correspond to one trial duration. Each trial involves four phases of functioning of the architecture: the planning phase, (possibly) the exploration phase, the action execution phase, and the learning phase.

At the beginning of each trial, and of the planning phase, the system observes a colour. During the planning phase, only the input units encoding the colour are activated, while the actions and feedback units are activated in a later phase. During each trial, a variable number of planning cycles is performed. A planning cycle represents the internal simulation of the trial events (colour, action, ation-outcome). The input layer is activated with the observed colour for a certain portion of the planning cycle (here 1/3 of its duration lasting 15 steps: for simplicity one planning cycle is assumed to last a number of steps as the actual trial, as done in [24]).

During one planning cycle, the arbitration mechanism operates as follows. The sequence sampling causes a certain activation of the neurons of the associative layer and a certain entropy averaged over the sampling steps. This entropy is considered a measure of the uncertainty of the world model. If this uncertainty is higher than a threshold, the arbitration component stops planning as not enough confident on the knowledge of the world model. Instead, if the uncertainty is lower than the threshold the arbitration component checks if the sampled sequence produced a state (read out in the output layer) that matched the goal, and if this is the case it stops planning and performs the action in the environment. Instead, if the arbitration component is confident on the world model but the sampling did not produce a sequence that matched the goal, it performs two operations before starting a new planning cycle. First, it updates the goal-associative connections so as to lower the goal-conditioned probability of the wrong sampled sequence. Second, it lowers the entropy threshold of a certain amount, thus ensuring that with time the probability of terminating the planning processes increases and the system does not get stuck in planning.

After planning terminates, if the system has not found an action that leads to a goal matching, the action is produced by the exploration component. The action selected either by the planning process or by the exploration component is performed in the environment.

#### Algorithm 1 Pseudo-code of the model functioning

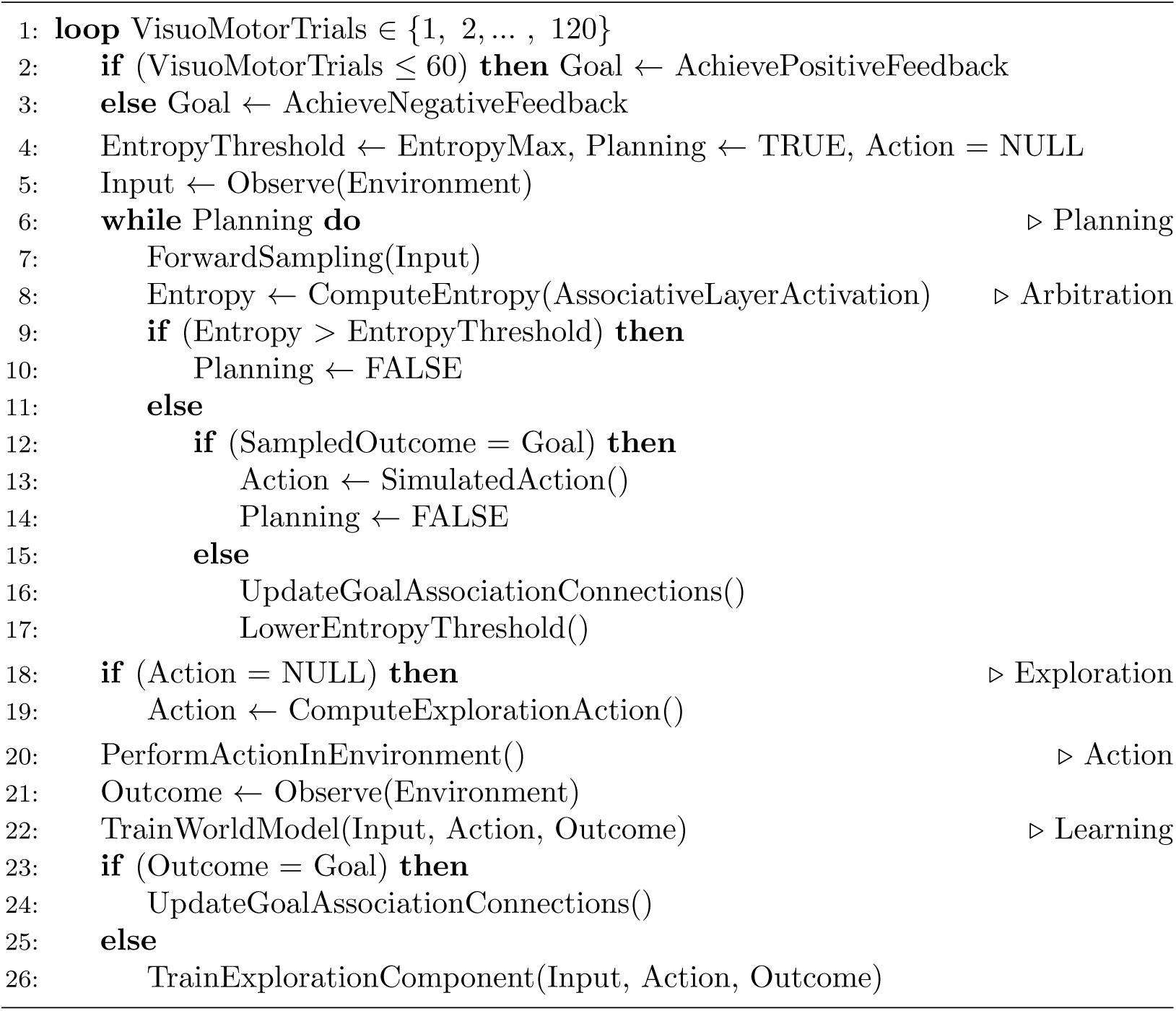

After action execution, the world model observes the feedback from the environment and also the action that caused it. Based on the observation of the stimulus (colour), action (finger press), and action-outcome (positive or negative feedback) from the environment, the world model learns. In particular, it learns the internal representation (hidden causes) of the observations (input-association connections), the possible time dependencies between them (association-layer internal connections), and the generation of the observations (association-output connections). Moreover, if the action led to actually reach the goal it increases the goal-conditioned probability of the sampled successful sequence (goal-associative connections). Instead, if the action failed only the exploration component is trained to avoid to produce the performed action in the experienced input-goal condition.

Note that when a trial starts, the architecture performs a planning cycle to evaluate entropy: this hypothesis is based on the fact that the task is novel. In a more general case where tasks could be familiar, a common habit/planning arbitration process might evaluate if a habit is available before triggering planning and the planning/exploration arbitration process considered here.

Note also that in case of goal-failure the goal-associative connections are updated during planning to exclude the multiple sampling of the same wrong sequence and action; instead, in the case of goal-achievement such connections are updated after the action is successfully performed in the world, rather than during planning, to avoid a training based on the possible false-positive errors of planning (false-negative errors are less likely during planning as the world model learns on the basis of the ground-truth information from the world). The exploration component is instead trained after the failure of the action executed in the world to avoid multiple explorations of actions found to be wrong (this hypothesis is inspired by the ‘inhibition-of-return’ mechanism of visual exploration, leading to exclude from exploration already explored items [39]); the component is instead not trained in case of success as this would amount to habitual learning not possible in few trials. These hypotheses were isolated through the search of the conditions for the correct reproduction of the target human data of the visuomotor task while fulfilling the challenging constraint that planning has to take place while learning the neural world model.

Based on these mechanisms, at the beginning of the visuomotor test the model tends to sample random stimulus-action-outcome sequences because the world model has no knowledge. The arbitration component thus quickly passes the control to the exploration component which decides which action to execute, and this is performed in the environment. With the accumulation of experience trials, the world model improves by learning the hidden causes of observations (colours, actions, feedback) and the time dependencies between them. This leads the arbitration component to measure a higher confidence in the world model, so planning continues and samples with a higher probability (hidden causes of) colour-action-feedback sequences that actually exist in the world. When one of these sequences leads to an action that predicts a goal achievement in the output layer, and the action is actually successful when performed in the environment, this leads to increase the goal-conditioned probability of sampling such sequence so that the next time the same colour is encountered the sequence is readily selected by the planning process.

### 1.3 Goal-directed behaviour model: detailed functioning

#### 1.3.1 The hidden Markov model represented by the world model

In this section we illustrate the aspects of a probabilistic HMM that are implemented by the spiking neural network world model. The HMM considers the hidden causes of world states, *h*_*t*_, and observations of them, *o*_*t*_, as random variables at the time steps *t* ∈ {0, 1, …, *T*} forming the sequences *H* = {*h*_0_, *h*_1_, …, *h*_*T*_} and *O* = {*o*_0_, *o*_1_, …, *o*_*T*_} The joint probability of these sequences can be expressed and factorised as follows given the assumptions on the probability independencies of the model shown in Fig 1A:

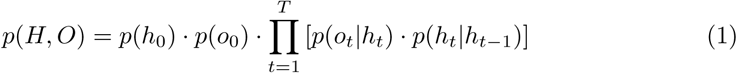

This formula highlights the two key elements of the HMM, namely the *generative model* of how the world states (hidden causes) cause the observations, *p*(*o*_*t*_|*h*_*t*_), and the *prediction model* of how a world state causes the following state *p*(*h*_*t*_|*h*_*t*−1_) (in the neural implementation of the HMM we will equivalently consider *p*(*h*_*t*_|*o*_*t*−1_) and *p*(*o*_*t*_|*h*_*t*−1_) to follow the general rule of the dependency of the state of any part of the neural network from the state of other neural parts at the previous time step).

The HMM has parameters *θ* that are adjusted on the basis of collected data (observations) so that the probability distribution *p*(*O* |*θ*) converges towards the empirical distribution from the world, *P* ^*^(*O*):

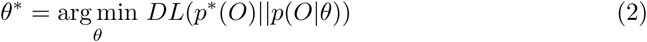

where *DL*(·‖·) is the Kullback-Leibler divergence between the two distributions and *θ*^*^ are the searched optimal parameter values. This problem cannot be solved in a close form and so *θ*^*^ are commonly searched numerically, in particular through an *expectation-maximisation* (EM) algorithm. Here we refer to how this is done in stochastic versions of HMMs [24, 40], most similar to the neural implementation of the HMM considered here. For these problems, the EM algorithm converges towards the solution by alternating an estimation step (E-step) and a maximisation step (M-step): the E-step samples a sequence of hidden causes *H*′ based on the posterior distribution *p*(*H*|*O*′, *θ*) dependent on the actual observations (*O*′); the M-step adjusts *θ* to increase *p*(*H*′|*O*′, *θ*). In the E-step, the sampling of *H*′ given *O*′ can be approximated by *forward sampling* [41], i.e. by sampling the *h*_*t*_ distributions in sequence, staring from *h*_0_, given the 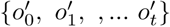 values observed until *t*.

#### 1.3.2 The spiking neural-network world model

The neural implementation of the world model instantiating the HMM is based on two learning processes. The first learning process, involving the input-associative connections, expresses and learns the hidden causes of observations as probability distributions of the spikes of the neurons of the association component at different time steps. The second learning process, involving the connections internal to the associative component, expresses and learns the temporal dependencies between the hidden causes of observations, reflecting the temporal dependencies between the states of the world, as conditional probability distributions of the spikes of the neurons of the association component at succeeding time steps.

The membrane potential of each neuron of the associative layer reflects the activation that would result from the typical connectivity pattern of cortex and other areas of brain, formed by neurons that reciprocally inhibit through inhibitory interneurons. This connectivity pattern tends to keep constant the overall firing rate of the layer. More in detail, the membrane potential *u*_*k*_ of a neuron of the model is:

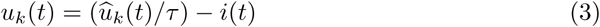

where *τ* is a scaling factor, *i*(*t*) is the common inhibition received by all neurons caused by the inhibitory interneurons to which they project, and 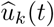 is the total activation received from other neurons:

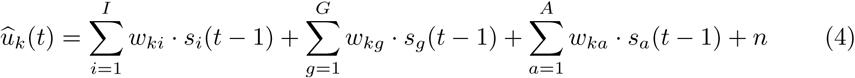

where *w*_*ki*_ are the input-associative connection weights, *w*_*kg*_ are the goal-associative connection weights, *w*_*ka*_ are the internal associative connection weights, *s*_*i*_(*t*), *s*_*g*_(*t*), and *s*_*a*_(*t*) are the incoming spike signals (*s* ∈ 0, 1) from the neurons of respectively the input, goal, and associative layer, and *n* is a Gaussian noise component with a standard deviation *ν*.

We then assume, as in [42], that the firing rate *v*_*k*_(*t*) of a neuron *k*, reflecting its spiking probability, is exponentially dependent on the membrane potential:

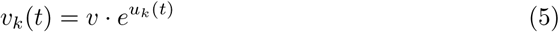

where *v* is a constant scaling the firing rate. This implies the following dependency of the neuron firing rate on the activation received by all neurons of the layer:

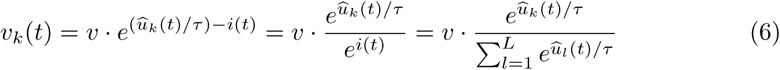

where *i*(*t*) was assumed to be:

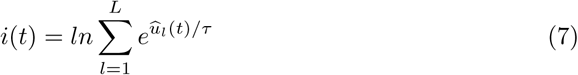

While the model on which we built assumed a continuous time and an inhomogeneous Poisson process to produce the actual spikes of the layer [24], we considered a discrete time, a fully Markov dependence between succeeding events, and a constant firing rate at each time step, assumed to be *v* = 1 without loss of generality. These assumptions simplified the analysis of the system and did not alter the core functioning of the model, in particular with respect to the effects of the core unsupervised learning rule illustrated below. With these assumption, Eq 6 becomes a *soft-max* function where 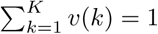 is the layer constant total firing, and *v*(*t*) can be interpreted as *v*(*t*) = *p*_*t*_(*k*), with *p*_*t*_(*k*) being a categorical probability distribution indicating the likelihood that the neuron with index *k* is the one to fire a spike at time *t* while the other neurons remain silent. The neurons of the output layer, receiving afferent connections from the associative layer, have the same activation as the neurons of the associative layer.

The weights of the connections linking the input-associative layers, the associative neurons between them, and the associative-output layers are updated through a Spike-Timing Dependent Plasticity (STDP) rule [43–46]. In particular, we used the following STDP learning rules from [24, 35] to update a connection weight *w*_*post,pre*_ linking the pre-synaptic neuron *pre* to the post-synaptic neuron *post*:

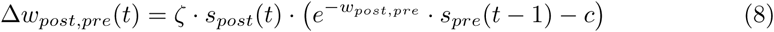

where *ζ* is a learning rate parameter, Δ*w*_*post,pre*_ is the size of the connection weight update, *s*_*pre*_(*t* − 1) and *s*_*post*_(*t*) is the spike activation (*s* ∈ {0, 1}) of respectively the pre-synaptic neuron in the current time step and post-synaptic neuron in the last time step, and *c* is a constant (*c* ∈ [0, 1]). The formula operates as follows. The rule updates the weight only when the post-synaptic neuron fires (*s*_*post*_(*t*) = 1). When this happens, but the pre-synaptic neuron does not fire (*s*_*pre*_(*t* − 1) = 0), then *w*_*post,pre*_ decreases of − *ζ* · *c*. This leads the post-synaptic neuron to form negative connections with all the pre-synaptic neurons that tend to not fire before it fires. Instead, if the pre-synaptic neuron fires before the post-synaptic neuron (*s*_*pre*_(*t* − 1) = 1), then *w*_*post,pre*_ increases if 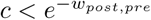 and decreases otherwise. This implies that *w*_*post,pre*_ will tend to converge to the positive attractor point 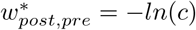 reached when 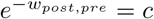. Overall, for a given neuron the rule tends to form positive incoming connections from neurons that fire just before it fires, and negative connections from all other neurons.

The connections that form are the means through which the system implements conditional probabilities. For example, initially the associative units *k*, each representing possible hidden causes of observations, tend to fire with a certain prior probability distribution, say *p*(*k*). The formation of input-associative connections allows an observation *i* to generate the posterior conditional probability distribution *p*(*k*|*i*) that for example implies an increased probability of selection of the hidden cause *k*.

Within the associative network, the learning rule leads to form a connectivity that supports a sequential activation of the neurons encoding the hidden causes of the observations, where the sequence reflects the temporal order in which the observations, reflecting the world states, are experienced by the HMM. The reason is that once the hidden causes are formed, based on the input-associative connections, then they tend to fire in sequence under the drive of the observations. As a consequence, the learning rule leads each associative neuron to connect with the associative neurons that fired before it and to form negative connections with those that did not fire. In this way, the connections within the associative network tend to form chain-like neural assemblies. These connections are hence able to represent the temporal dependencies between hidden causes, e.g. between *a* and *k* corresponding to two succeeding observations, as conditional probabilities *p*(*k*|*a*). Importantly, if the system observes different events following the initial observation of the trial (e.g., different actions and different outcomes after a certain initial colour), the world model will exploit its stochastic neural processes to represent such possible alternative sequences of events. This is at the core of the architecture’s capacity to internally simulate alternative courses of actions and events and hence to plan in a goal-directed manner.

The same learning rule is also used to train the associative-output connections. Initially, the output layer expresses a probability distribution, say *p*(*o*), that tends to be uniform and so when sampled generates unstructured observations. With learning, the world model strengthens some connections between the spiking sequences sampled within the associative network and the observations activating the output layer. When the world model samples an internal sequence within the associative network, this leads to generate the observations on the basis of the reconstruction probability *p*(*o*|*k*).

Overall, the neural HMM plus the output layer act as an *auto-encoder* returning as output the input, and able to capture in its internal states the hidden causes of observations: this is similar to what happens in variational auto-encoder [47], a probabilistic version of auto-encoder [48], with the difference that the model considered here generates sequences of patterns rather than single patterns. In this respect, the output layer acts similarly to the reading-out layer of a dynamic reservoir network [49, 50] which is however deterministic.

When the planning process has to generate an action to perform, or a predicted feedback to compare with the goal, the generated event at the output layer is considered to be the one that fired the most during the planning cycle. If the system had to generate sequences of events involving multiple actions and predicted states, one should consider other ‘reading out’ mechanisms, for example that an event is generated each time the unit encoding it fires a minimum number of spikes in sequence.

The goal-associative connection weights are updated on the basis of the success or failure to achieve the goal during planning or when the action is performed in the environment. In particular, the update is done on the basis of the following reinforcement learning rule:

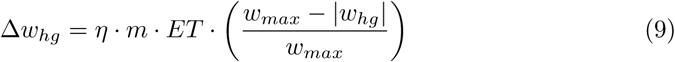

where *η* represents the learning rate, *m* is the reward, equal to 1 if the sequence resulted in a successful goal matching and −1 otherwise, *ET* is the Eligibility Trace, equal to 1 for units that have fired at least once during the planning cycle/trial and to 0 otherwise, and *w*_*max*_ is the maximum absolute value that the weight can reach (*w*_*max*_ = 0.5). The goal-associative connections allow the goal *g* to condition the probability distribution over the hidden causes, *p*(*k*|*i, a, g*). With learning, this allows the goal to condition the probability of the sampled hidden causes sequences so as to increase the likelihood of those that involve the correct action. Moreover, when the goal changes, the system is able to modify the conditioned probability of the sequences so as to increase the probability of sampling a new sequence, based on the same world model, achieving the new desired goal.

### 1.3.3 Arbitration system

The arbitration component decides if continuing to plan or to pass the control to the exploration component and/or perform the action selected by either the planning or the exploration process. The component pivots these decisions on a key information, namely an estimation of the level of knowledge of the world model for the given trial initial observation (here the colour). This knowledge is related to the fact that the world model has possibly learned some sequences of events (action-feedback) that might follow the initial observation. A good level of knowledge means that the probability mass of the distribution *p*_*t*_(*k*|*i, a, g*) during the planning cycle steps *t* is concentrated on few possible hidden causes. The measure of this knowledge is in particular based on the entropy of the probability distribution at each time step *t*:

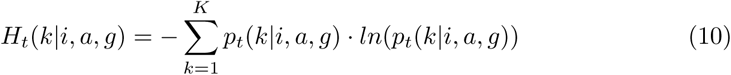

and the maximum value of such entropy is *H*_*max*_ = *ln*(*K*) corresponding to a uniform probability distribution where each *k* neuron of the layer has the same probability of firing *p*(*k*) = 1*/K*. The measure of the uncertainty *H* of the world model in a given planning cycle lasting *T* time steps is in particular defined as:

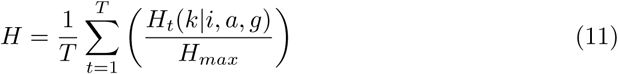

At the end of each planning cycle, the arbitration component computes *H*, compares it with an entropy threshold *H*_*Th*_(*t*), compares the action-outcome *z* with the pursued *g*, and selects one of three possible functioning modes of the architecture:

- *H* < *H*_*Th*_(*t*) and *z* ≠ *g*. The goal-associative connections are updated and a new planning cycle starts.
- *H* < *H*_*Th*_(*t*) and *z* = *g*. Planning stops and the action of the last planning cycle that caused the anticipation of the goal is executed in the world (without activating the exploration component).
- *H*_*Th*_(*t*) < *H*. Planning stops and control is passed to the exploration component.

The entropy threshold decreases linearly at each planning cycle so that the exploration component is eventually called if the planning process fails to match the goal within a certain time:

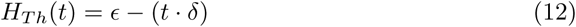

where *ϵ* is the value to which the entropy threshold is set at the beginning of the trial (and the planning process), and *δ* is its linear decrease.

The exploration component is a neural network formed by two layers. The first is an input layer formed by 6 neurons encoding the elements of the Cartesian product between the possible three colours and two goals. The second is an output layer formed by 5 neurons representing the possible actions, receiving all-to-all connections from the input layer. When the exploration component is called to select the action, the input layer is activated according to the current colour-goal combination (hot-vector activation), the activation potential of the second layer units is computed as usual as the sum of the weighed inputs, and an action is chosen on the basis of a *soft-max* function (Eq 6). When the action leads to a negative reward, the connection weights of the component are updated using the same reinforcement learning rule used for the goal layer (Eq 9). This leads to exclude actions that are not useful for the current state-goal combination, thus fostering exploration. Note that an additional slow-learning component similar to the exploration component might be used to model the formation of habits in experiments involving long learning periods.

### 1.4 Search of the model parameters

The model functioning depends on seven important parameters, indicated in Tab 1. We searched the best values of those parameters by fitting the model behaviour to the corresponding data of the human participants. In particular, we randomly sampled and evaluated 100,000 parameter combinations. For each combination, we recorded and averaged the behaviour of 20 ‘simulated participants’, in particular their performance in the 20 trials with the stimuli S1, S2, and S3, and the average reaction times over colours on the same trials (this because the original data on the reaction times of humans were not separated). Such three performance datasets and one reaction-time dataset were compared with the corresponding average data from 14 human participants through a Pearson correlation coefficient *R*_*d,m*_ computed as:

**Table 1.** Parameters identified with the grid search technique. In particular, parameter names, minimum and maximum range, and values found by the search.

| Name | Range min | Range max | Found value |
| --- | --- | --- | --- |
| STDP learning rate ($\zeta$) | 0.1 | 1.0 | 0.96 |
| STDP threshold ($c$) | 0.1 | 1.0 | 0.67 |
| Planner learning rate ($\eta$) | 0.001 | 1.0 | 0.008 |
| Softmax temperature ($\tau$) | 0.01 | 0.1 | 0.02 |
| Neural noise ($\nu$) | 0.01 | 0.1 | 0.02 |
| Entropy max threshold ($\epsilon$) | 0.3 | 1.0 | 0.74 |
| Entropy decrease ($\delta$) | 0.01 | 0.2 | 0.12 |

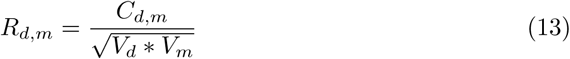

where *C*_*d,m*_ is the covariance between the data from humans, *d*, and data from the model, *m*; *V*_*d*_ and *V*_*m*_ is their respective variance. In particular, the coefficient was computed separately for the different data sets (performances and reaction times) and then averaged.

The range of the parameters explored by the search, and the best parameter values that it found, are shown in Tab 1. The best parameter values, that had a correlation coefficient of 0.72, were used in all the simulations illustrated here.

## 2 Results and Discussion

This section illustrates the behaviour and functioning of the model when tested with the visuomotor learning task proposed in [12] and described in Sec 1.1. The reported results refer to twenty replications of the simulations each representing a simulated participant performing the task. The results are also discussed from the perspective of the current state-of-the art on probabilistic and spiking neural-network models of goal-directed behaviour.

### 2.1 Behavioural analysis

Fig 4 shows that the model exhibits a performance similar to the human participants. The performance is in particular very similar to the humans’ one for the colours whose correct action is found after one or three errors whereas it is slightly lower for the colour whose correct action is found after four errors.

**Fig 4.**
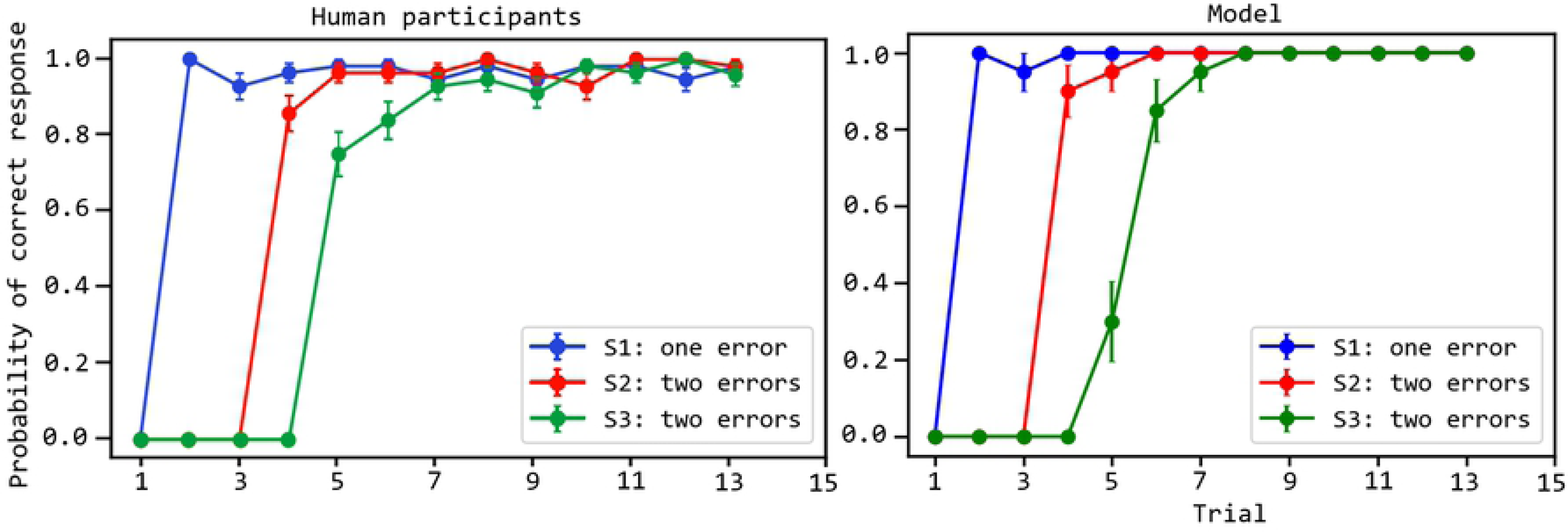
Comparison of the performance of the human and simulated participants. The performance (y-axis) is measured as the proportion of correct response over the trials (x-axis), separately for the three different colour stimuli (S1, S2, S3). Curves indicate the values averaged over 14 human participants and 20 artificial participants; error bars indicate the standard error. The data of human participants are from [38].

Once the model finds the correct action for one colour, when it encounters the same colour again it reproduces the correct action with a high probability. The architecture however takes more cycles to converge to such a high probability for S3 because the planner has a larger number of wrong sequences and so has a higher probability of wrongly anticipating a positive feedback. This problem is less impairing for S1, and in part for S2, involving fewer wrong sequences during planning.

The arbitration component decides to implement a different number of planning cycles (each involving the generation of colour-action-feedback sequences) depending on the knowledge stored in the world model. If a larger number of planning cycles is performed, the reaction times of the architecture are considered to be longer. These reaction times can be compared with those of the human participants (Fig 5). The reproduction of the human reaction times is particularly interesting and challenging as it has an inverted ‘U’ shape.

**Fig 5.**
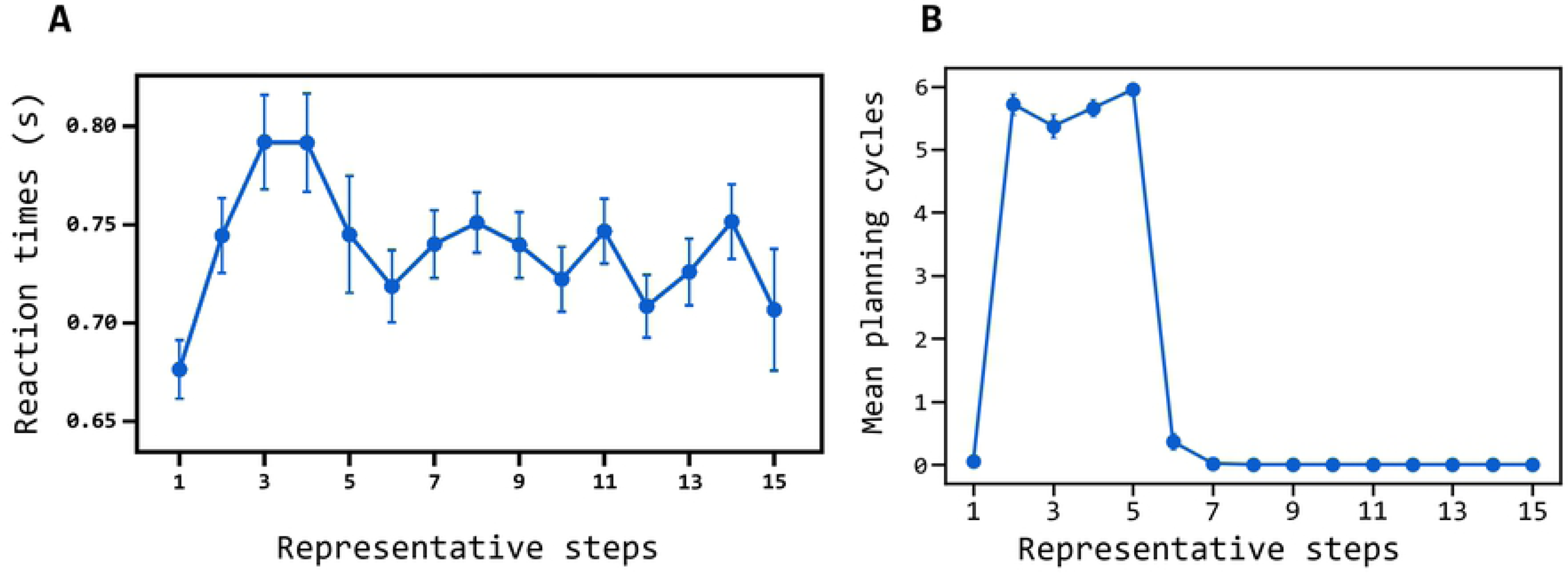
Comparison of the reaction times of the humans and simulated participants. (A) Reaction times of human participants averaged over S1, S2, and S3 (y-axis) for the ‘representative steps’ (x-axis); the ‘representative steps’ allow the alignment of the reaction times of the three stimuli so as to separate the exploration phase (first 5 steps) and the exploitation phase (6 steps onward); to this purpose, the reaction times for S1 obtained in succeeding trials from the first onward is assigned the steps (used to compute the averages shown in the plot) ‘1, 2, 6, 7, …’, whereas S2 is assigned the steps ‘1, 2, 3, 4, 6, 7, …’, and S3 is assigned the steps ‘1, 2, 3, 4, 5, 6, 7, …’; data are taken from [38]; (B) Reaction times of the model plotted in the same way.

In the first trials, for each stimulus the entropy (uncertainty) of the world model is high as the associative layer expresses a rather uniform probability distribution. Indeed, the component has still to identify the hidden causes of stimuli and actions, so the neurons forming it tend to spike with a similar rate. As the entropy is high, the arbitration component tends to quickly pass the control to the exploration component and so the reaction times are low. After the following trials in the environment, the world model start to form representations of the experienced colour-action-feedback sequences and to assigning to them a higher posterior probability with respect to other patterns. The arbitration component thus tends to compute a lower entropy, the architecture plans for longer, and so the reaction times get longer. During this planning, the associative component tends to sample the learned sequences with a high probability conditioned to the observed colour. If none of the sequences leads to predict an event that matches the pursued goal through the output layer, the probability of such sequences is however decreased under the conditioning of the goal and the control is again passed to the exploration component. When the action performed in the world manages to produce the desired goal, the system learns the corresponding sequence and assigns to it a high posterior probability. When the colour of such sequence is observed again, the sequence is sampled with a higher probability and results in a successful match. The arbitration component stops planning and the action is performed in the world. The reaction times are hence short again.

In summary, the inverted ‘U’ shape of the reaction times is caused by these processes: (a) initially the world model has learned no sequences, entropy is high and overcomes the threshold, and so the arbitration component passes the control to the exploration component: the reaction times are low; (b) when the world model has learned some sequences but these are wrong, planning implements several cycles to explore such sequences and to lower their goal-conditioned probability, so the arbitration component takes time to pass the control to the exploration component: the reaction times are high; (c) when the world model has learned the correct sequence, entropy is low but the planning process samples such sequence with a high probability, obtains a successful matching of the goal, and the found successful action is performed: the exploitation of the found solution starts and the reaction times become low again.

The results on the performance and reaction times of the model allow us to discuss two of the key features of the model with respect to the existing literature about probabilistic planning models based on spiking neural networks. The first feature is that the world model is learnt in parallel with its use for planning, and an arbitration mechanism decides when to explore or to plan on the basis of an entropy-based confidence on the world model. Previous models of probabilistic planning based on spiking neural networks did not consider the possibility of using approximate world models (as these were trained before the solution of tasks) and a mechanism of arbitration to decide if planning or exploring [35, 36]. One of the first models that studied arbitration [37], but that did not rely on spiking neural network, learned the dynamics of the world as a *Bayesian decision tree* for planning and in parallel also learned habitual behaviour based on reinforcement learning (*Q-learning*). The arbitration system decided if using planning or habitual behaviour on the basis of the variance of the state-action values of the two components. Since the model used here is grounded on neural computations, we could compute the confidence on the world model through a measure directly linked to those neural computations, namely the entropy of the probability distributions of the world-model neural network spikes. In future work, this could allow the computation of such measure through a biologically plausible neural mechanism.

Another model [38] used an entropy-based measure as a means to decide to give control to a goal-directed component or to a habitual component. Here the goal-directed component was based on a *Bayesian Working Memory* (a memory of the one-step time-dependent state probabilities, and of the one-step transition-function and reward-function probabilities) and the habitual behaviour was based on Q-learning. The model was validated with the same visuomotor task used here. The model reproduced the reaction times of the target experiment by making them dependent on the sum of two elements: (a) the logarithm of the number of items in working-memory, corresponding to the performed trials (this affected reaction times only when planning); (b) the entropy of the action probabilities. The inverted ‘U’ shape of reaction times was produced by the fact that the first component of reaction times tended to increase with experience accumulating items in working memory, and the second component tended to decrease with the decrease of the variance of the action probability distribution. Instead, here the inverted ‘U’ shape is an emergent effect of the change of knowledge of the world model traversing three phases: (a) no knowledge: fast reaction times; (b) ample knowledge on the world functioning, but little knowledge on which part of it is relevant for the goal: slow reaction times; (c) ample knowledge on the world functioning, and specific knowledge on which part of it should be used to accomplish the goal: fast reaction times. The empirical and computational implications of the two different hypotheses deserve further study.

Another difference of our model with respect to the models of [37, 38] is that it uses arbitration to select between exploration (attempts of different actions serving the world-model learning) vs. exploitation of the acquired knowledge (planning), rather than between goal-directed behaviour and habitual behaviour. This was done as habitual behaviour takes long to form and so it seems to be ruled out during the first attempts to solve new problems [3]. Instead, the first phase of solution of a new task involves the learning of the model of the world based on the exploration of how the world responds to actions, and the possible exploitation of the collected knowledge by the goal-directed components. In this respect, the arbitration component proposed here makes decisions at a finer time granularity with respect to other models, namely during planning cycles, rather than at the coarser granularity of the trials, as in the other models. Future work could investigate how to add to the exploration/exploitation processes used here the contribution of an habitual component that would slowly form with experience. This component might have an architecture similar to the exploration component, and learn on the basis of positive reward from experience in the world. This would require an interesting integration of the exploration/exploitation arbitration mechanisms used here and the goal-directed/habitual arbitration mechanism used in previous models.

Another difference linked to the previous point is that the model presented here used an explicit representation of the *goal* to directly (learn to) condition the probability distributions of the world model, rather than to generate a reward corresponding to the desired state and used it to perform reinforcement learning based on the world model (model-based reinforcement learning) as in [37, 38]. Our different approach was also uses in [35] who however conflated the goal, initial state, and environment conditions into a whole ‘context’ representation. Instead, in the model presented here the initial state and goal have factored representations, and the ‘environment condition’ is considered as part of the world model state. Thus the model is able to express the goal-free probability distributions representing the ‘objective dynamics’ of the world, or to express the ‘goal-based probability distributions’ due to the agent’s action to accomplish the goal. The implications of this shift of perspective deserves further study, in particular to infer possibly different empirical predictions and understand the possible computational advantages/disadvantages.

### 2.2 Model dynamics

Fig 6 shows how the activation of the associative layer evolves during the planning trials, across the succeeding trials of the test due to the increased knowledge of the world model (recall that planning might involve a different number of forward samplings). Initially (trials T1-T3), the prior probability of activation of the neurons of the associative layer tends to be uniform, thus resulting in a random sampling (spiking) of the neurons. This means that the model has still not identified specific hidden causes of the observations.

**Fig 6.**
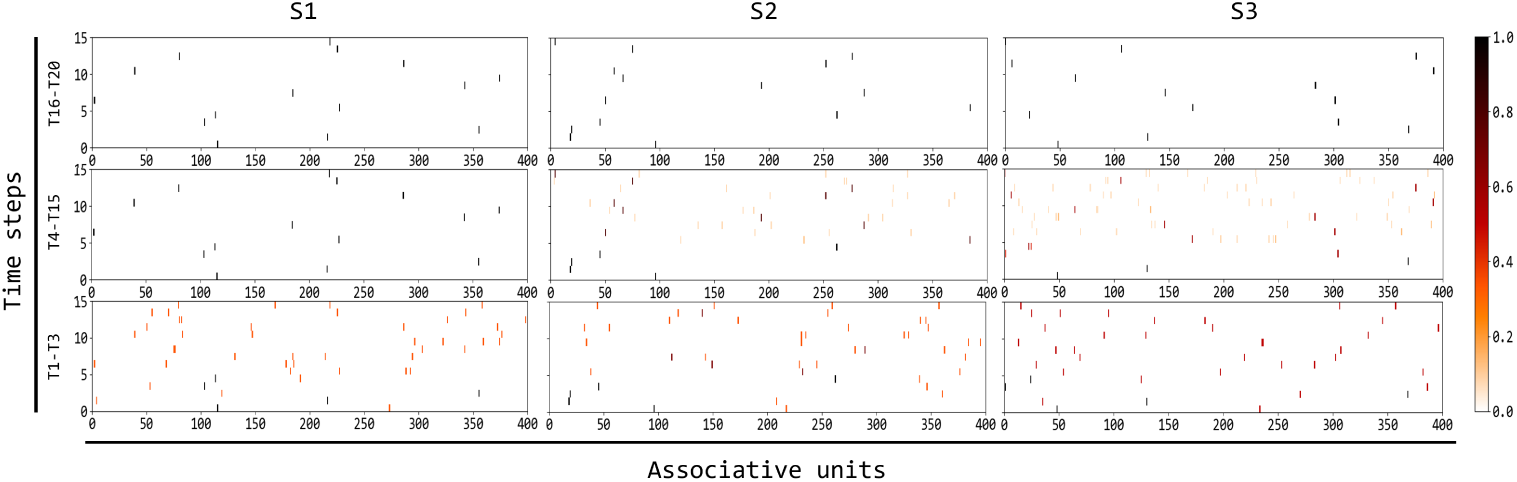
Evolution of the spiking activity of the associative units during planning. The three rows of graphs correspond to different succeeding sets of trials (T1-T3, T4-T15, T16-T20) used to compute the average activation of the units shown in the graphs (the colour indicates such average, normalised in [0, 1]), and the three columns of graphs refer to the three possible colour stimuli (S1, S2, S3). Each graph shows the average activation of the 400 units of the associative network (x-axis) during the planning forward samples (y-axis). Recall that each trial involves: the perception of the stimulus colour, planning (involving a different number of forward simulations of the trial events: the graphs show the firing of the associative neurons during these sequences), (possibly) the activation of the exploration component, and finally the performance of the action in the world.

With the experience of the input stimuli, the STDP acting on the input-associative connections and on the internal associative connections leads the associative layer to form an internal representation of the hidden causes of the observations, namely of the colour, the action, the feedback, and the elapse of time (the latter due to the fact that each observation lasts multiple time steps). At the same time, the plasticity of the associative layer leads it to form a HMM that represents in an increasingly accurate fashion the time-related probabilistic dependencies between the discovered hidden causes. Finally, once some possible sequences are encoded by the associative component starting from the current colour, the STDP acting on the goal-associative connections progressively increase the probability of sampling sequences that lead to achieve the goal and to decrease the probability of those that do not. The effect of these processes can be seen in the figure graphs, in particular with respect to S1 for which a successful sequence is discovered after two trials (three graphs at the left). For this stimulus, during trials T4-T15 some specific neurons start to fire in sequence more strongly than other neurons, meaning that the system has learnt to represent the hidden causes, and their time dependencies, of the events of the first successful colour-action-feedback sequence.

During T4-T15 and T16-T20, the world model also learns the hidden causes, and their temporal dependencies, of the events of the other two sequences corresponding to S2 and S3 (second and third column of graphs in the figure). Here, learning of the world model and its correct exploitation takes more trials with respect to S1 as the goal (successful feedback) is achieved after a larger number of sequences (four and five for respectively S2 and S3). This implies that the architecture takes longer to first learn the hidden causes of all such sequences and then to decrease the probability of the wrong ones based on the pursued goal.

Importantly, during these experiences the world model, which tends to record any aspects of the world dynamics independently of the fact that it is useful to pursue the currently goal or not, also learns sequences leading to a negative feedback. The next section shows how this knowledge might become useful to accomplish other goals.

Overall, the figure shows that the operation of STDP leads to these effects: (a) the system’s world model learns to represent the elements and time relations of the observed sequences of events independently of the pursued goal; (b) when event sequences are learned, they can be sampled by the planning process to figure out the actions to use to pursue the desired goal; (c) the successful achievement of the goal later leads, when the goal is active, to increase the probability of sampling the sequences of events involving the correct actions.

Fig 7 shows, with analogous graphs, how the activation of the output layer during the planning trials evolves in time due to the increasing knowledge acquired by the world model. The firing of the output layer during planning expresses the predictions of the events that might happen starting from the observed trial colour. Such predictions are based on the simulation of the possible evolution of the world events based on the HMM instantiated by the associative layer. Regarding S1 (left three graphs of the figure), during the first trials (T1-T3) the world model has no/little knowledge on the dynamics of the world, so the activation of the units in the output layer reflect a uniform probability distribution leading to random predictions of the trial events. With additional experiences of trials involving S1 (T4-T15), the world model starts to learn to represent the trial events and, under the conditioning of the current goal, to assign a high probability to the correct colour-action-feedback sequence. As a consequence, the probability distribution of the output layer starts to correctly predict such correct sequence.

**Fig 7.**
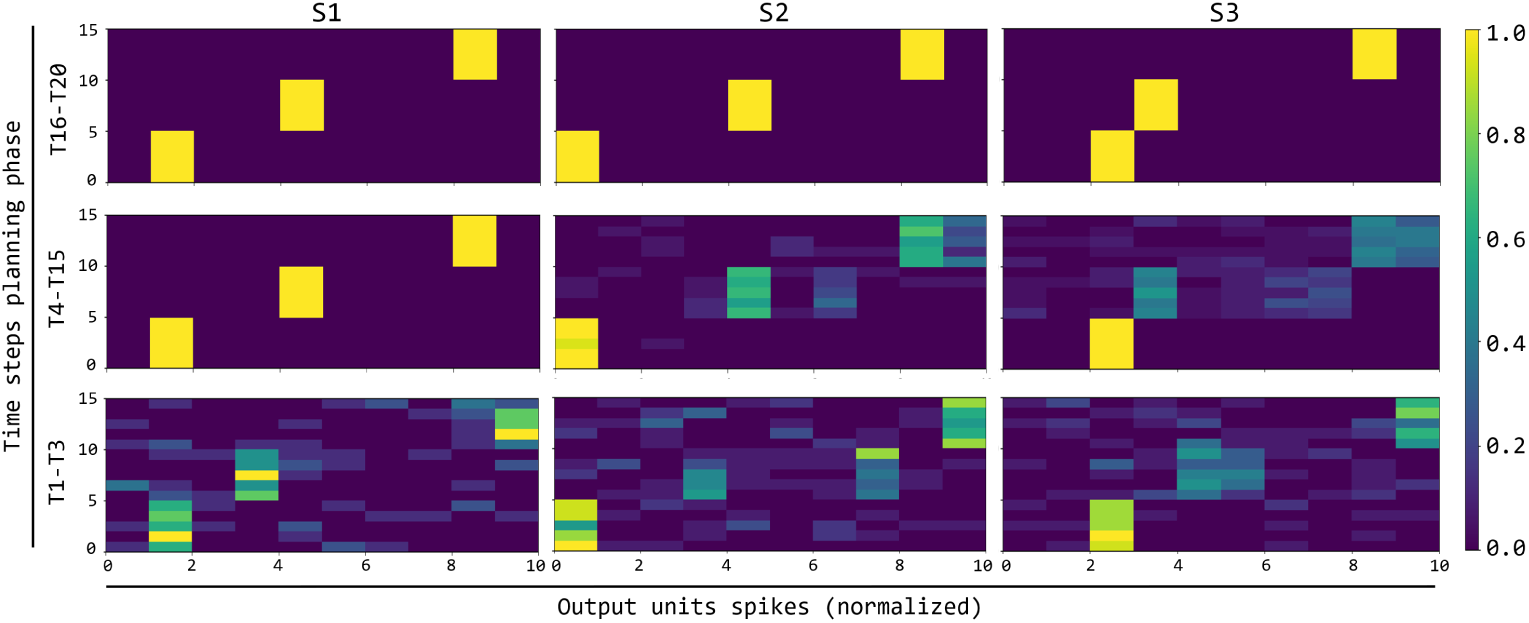
Evolution of the activation of the output layer units encoding the predicted observations and actions. The three rows of graphs correspond to different succeeding sets of trials (T1-T3, T4-T15, T16-T20) used to compute the average activation of the units shown in the graphs (the colour indicate such average, normalised in [0, 1]), and the three columns of graphs refer to the three colour stimuli. Each graph shows the activation of the 10 units (x-axis; units 1-3 encode the three colours, units 4-8 encode the 5 actions, and units 9-10 encode the positive/negative feedback) during the 15 steps of the trials (y-axis).

During trials T4-T15 and T16-T20 the same process happens for the correct sequences of the two colours S2 and S3. Also for these stimuli, towards the end of all trials (T16-T20) the probability distribution expressed by the output layer, conditioned to the associative layer activation, has converged to a probability close to 1 for the correct sequences.

These results allow the discussion of a novel feature of the model with respect to other systems implementing planning as inference based on spiking neural networks [35, 36]. In these systems, the world model considers possible sequences of states while abstracting over the actions that might lead to them: actions are computed ‘off-line’ with respect to the planning processes searching possible state sequences to the goal. Moreover, the world model is trained during a random exploration of the environment where actions are chosen according to a uniform probability distribution. As a consequence, the world model can only reflect this probability distribution: given a new goal, the system has thus to infer the probabilities of new possible sequence of states and actions from scratch. Instead, the world model used here is a HMM that observes states and actions as similar events, independently of the fact that they are produced by the environment or by another part of brain (e.g., the actions produced by the exploration component used here, or by a future habitual component). This allows the world model to learn state-transition probabilities that are sensitive to the probability of action selection. This might allow two possible advantages. First, it could support the biasing of the action selection probabilities, and hence the state probabilities, in favour of actions that lead to potentially useful effects from a given state, rather than any action in any state. This might be used to bias the world model to produce sequences of events involving only actions that are useful to accomplish states relevant for the agent’s typical goals (this might capture the important concept of *affordance* used in cognitive sciences [51], see [52]). Moreover, the use of state-action sequence probabilities might allow goals to bias only the probabilities of action elements of the HMM rather than also the probabilities of the state elements, that could thus usefully reflect the actual physics of the world. If a distributed representation of goals is used that allows generalisation over them, this would for example allow new goals similar to previously goals to immediately bias the action probability distribution, and hence the state probability distribution, expressed by the HMM in favour of potentially relevant actions and states.

A further novelty of the model presented here with respect to neural probabilistic models [35, 36] is that the world model is more realistically learned during the solution of the novel task, rather than before the task solution. This caused the challenge of using a partial model of the world for planning, faced with the novel exploration/exploitation arbitration mechanism proposed here (this same challenge was faced by previous models, [37, 38], but these used goal-directed/habitual arbitration, and were not grounded on spiking neural networks).

Another novelty of the world model presented here is that it learns on the basis of a biologically-plausible *unsupervised* neural learning mechanism [24], rather than based on the indication of the internal desired activation patterns by an external ‘teacher’ [35, 36]. Computationally, finding the conditions for the successful functioning of such unsupervised learning process, together with the acquisition of the world model while using it for planning, represented the hardest challenge for the construction of the architecture proposed here.

### 2.3 Model empirical prediction

An important advantage of planning is that the world model can store general knowledge on the dynamics of the world that can be used to accomplish different goals. It was thus interesting to check to which extent the current architecture preserved this capacity since it incrementally acquires a partial world model while solving the visuomotor task. To this purpose, after the architecture underwent the experiences reported in the previous section, it was required to perform additional trials to pursue the different goal of ‘obtaining a negative feedback’ in correspondence to the three colours. As shown in Fig 8A, when the goal is switched, the architecture is able to quickly change behaviour and choose the sequences that lead to the desired new goal given the colour. What happens is indeed that, under the conditioning of the observed colour, the world model already represents the hidden causes of the elements of the sequences and also assign a high probability to these sequences. In particular, since the previous goal unit is now off, the probability of the different sequences tends to be similar, and so the system tends to sample all of them equally during planning. This allows the architecture to rapidly discover a sequence that leads to the desired new goal, to solve the new version of the task through it, and then to increase the probability of such sequence conditioned to the new goal.

**Fig 8.**
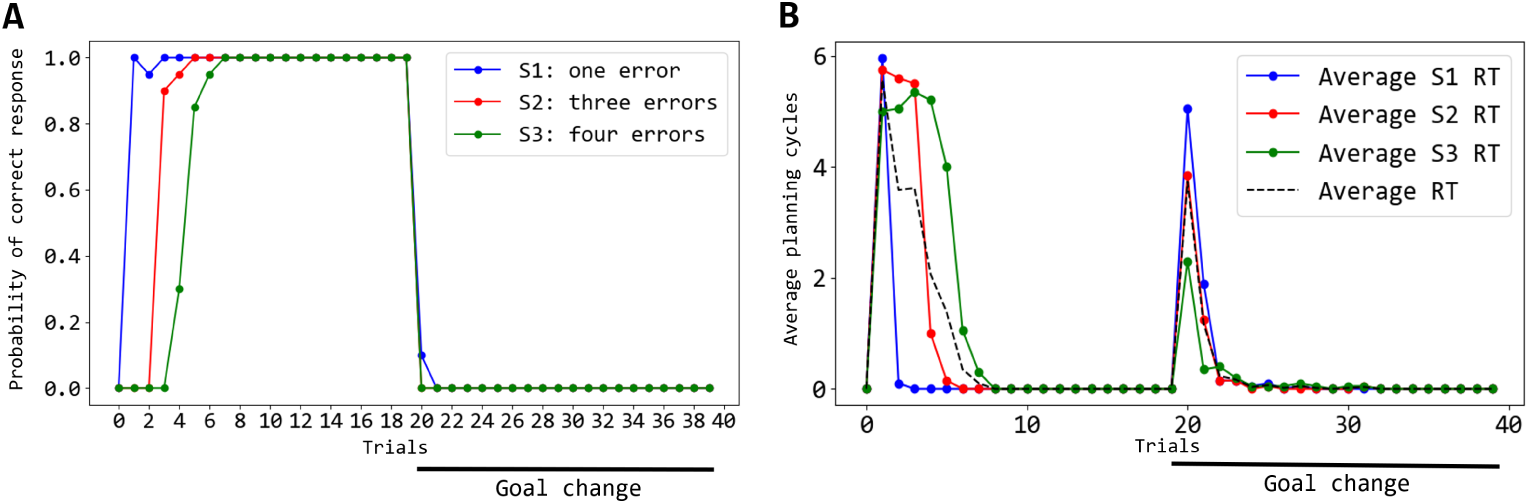
Behaviour of the system when the goal is switched to a new one, averaged over 20 simulated participants. (A) Performance measured as probability of selection of the ‘positive feedback’ goal: the pursued goal is switched from ‘getting a positive feedback’ to ‘getting a negative feedback’ at trial 20. (B) Average reaction times measured during the same experiment shown in ‘A’.

Regarding the reaction times (Fig 8B), the model shows a transient increase of their size in correspondence to switch of the goal. This is due to the fact that with the new goal the system needs to perform the sampling of some sequences before finding the successful ones. The reaction time is higher for S1 than for S2 and S3 as for it the model has less sequences available to reach the new ‘negative feedback’ goal and constantly one sequence to achieving the ‘positive feedback’.

These results represent a prediction of the model that might be tested in future experiments with human participants resembling the simulated test performed here (never performed with humans). In particular, the model predicts a certain performance and reaction times (Fig 8), possibly distinct for stimuli S1/S2/S3, that might be measured and compared with those of humans.

Overall, these results show how once the world model has acquired goal-independent knowledge on the environment dynamics the architecture can use it to pursue different goals. This feature is the hallmark of the flexibility of goal-directed behaviour and is shared with the other neural probabilistic models [34, 36]. These, however, were not validated with specific empirical data and were not used to produce specific empirical predictions as here.

## 3 Conclusions

Goal-directed processes represents a key process supporting flexible behaviour based on general-purpose knowledge of the world. In recent years, it has been proposed that planning processes are based on probabilistic representations of the world and inferences on them. This proposal is very interesting but it encounters the great challenge of explaining how such representations and processes might be grounded on the actual neural computations of the brain. Recently, some models have been proposed to ground some probability inference mechanisms, such as Hidden Markov Models and Partially Observable Markov Decision Processes, on the spiking stochastic events exhibited by the brain neurons, and on some typical brain connectivity patterns and plasticity mechanisms.

Here we contributed to bridge probabilistic planning to brain mechanisms by proposing a goal-directed model that implements planning based on probabilistic representations and inferences grounded on spiking neural networks. The model was tested with data on human participants engaged in solving a visuomotor behavioural test that requires to discover the correct actions to associate to some stimuli [12]. The model reproduced the target behaviour, furnished an explanation of the mechanisms possibly underlying it, and presented predictions testable in future empirical experiments.

The model has three novelties with respect to existing probabilistic planning systems, in particular with respect to models relying on brain-like mechanisms. First, the model is able to perform generative planning using a partial model of the world, as commonly happens when a new task is solved. The model can do this based on an arbitration mechanism that decides to plan if relevant knowledge has been acquired by the world model, or to explore the environment to acquire further knowledge. Second, the world model is based on a Hidden Markov Model that predict states as well as actions, and can bias the probability with which spikes sample such items on the basis of the pursued goals. Last, the learning of the hidden causes of observations and their temporal dependencies are learned through an unsupervised STDP mechanism that enhances the possibilities of linking the model to brain.

Various aspects of the model might be improved in future work. A first one concerns the passage from neurons firing at discrete times to neurons firing in continuous time. This might be done using the not homogeneous Poisson process used in [24]. Although this would not change the theoretical contribution of the model, it might simplify a comparison of the model functioning with real data from brain at a finer temporal level with respect to the one considered here.

A further issue to face would be to use other tasks with respect to the one considered here [12], so as to develop the model to tackle longer sequences of states and actions, e.g. as in [35, 36]. As done in these works, it would also be interesting to employ the model as the controller of autonomous robots to test its robustness and versatility when facing more complex tasks.

Another improvement might involve the full development of a habitual component. Here we did not introduce such component as the target data seemed to not require it, so we focused on considering the exploration/exploitation phases involved in the learning of the new task. Future work might also consider habits, e.g. by referring to additional target experiments involving a long ‘over-training’ of behaviour. The current exploration component of the model has already an architecture suitable for this. The addition of habitual processes would however also require a more sophisticated arbitration mechanism, e.g. by integrating ideas from [37, 38].

A further possible improvement of the model concerns the goals. These are now selected externally and represented in a simple way. Goals could instead be represented in more realistic ways, e.g. through mechanisms mimicking working memory [53], and could be selected in autonomous ways, e.g based on motivational mechanisms [54].

A last improvement concerns the possibility of testing and constraining the model not only at the behavioural level, as done here, but also at the neural level, for example based on data collected on the same experiment [55, 56]. This might for example be done through analysis techniques such as *Representational Similarity Analysis* [57] that maps the components of neural models to areas of brain possibly performing similar functions, on the basis of brain-imaging data.

Notwithstanding these possible improvements, the model represents a further step towards the development of an architecture implementing goal-directed behaviour that on one side is able to take into consideration the stochastic nature of the world, as in probabilistic planning, and on the other relies on the functioning and plasticity mechanisms that can be linked to those of brain.

## Acknowledgements

R.B.: Idea of the model, specification of the model and tests, implementation of the model, tests, data analysis, analysis of results, writing-up. G.B.: Idea of the model, specification of the model and tests, analysis of results, writing-up. E.C. and A.B.: Specification of the model and tests, analysis of results, writing-up. R.B. carried out this work thanks to the support of the A*MIDEX grant (n° ANR-11-IDEX-0001-02) funded by the French Government “Investissements d’Avenir” program (https://anr.fr/en/investments-for-the-future/investments-for-the-future/). G.B. and E.C. received funding from the European Union, Horizon 2020 Research and Innovation Program, under Grant Agreement n° 713010 Project “GOAL-Robots - Goal-based Open-ended Autonomous Learning Robots” (https://cordis.europa.eu/project/rcn/203543/factsheet/en). A.B. was supported by the French National Agency (n° ANR-18-CE28-0016-01) and the FLAG-ERA (n° ANR-17-HBPR-0001-02)(https://anr.fr/Projet-ANR-18-CE28-0016; https://anr.fr/Project-ANR-17-HBPR-0001).

## References

1. Dickinson A, Balleine B. Motivational control of goal-directed action. Animal Learning & Behavior. 1994;22(1):1–18. doi: 10.3758/BF03199951.

2. Balleine BW, Dickinson A. Goal-directed instrumental action: contingency and incentive learning and their cortical substrates. Neuropharmacology. 1998;37(4):407–419.

3. Dolan R, Dayan P. Goals and Habits in the Brain. Neuron. 2013;80(2):312–325. doi: 10.1016/j.neuron.2013.09.007.

4. Sutton RS, Barto AG. Reinforcement learning: an introduction. Cambridge, MA: The MIT Press; 1998.

5. Botvinick MM, Niv Y, Barto A. Hierarchically organized behavior and its neural foundations: A reinforcement-learning perspective. Cognition. 2008;113(3):262–280.

6. Balleine BW, Dezfouli A, Ito M, Doya K. Hierarchical control of goal-directed action in the cortical–basal ganglia network. Current Opinion in Behavioral Sciences. 2015;5:1–7. doi: 10.1016/j.cobeha.2015.06.001.

7. Mannella F, Gurney K, Baldassarre G. The nucleus accumbens as a nexus between values and goals in goal-directed behavior: a review and a new hypothesis. Frontiers in Behavioral Neuroscience. 2013;7. doi: 10.3389/fnbeh.2013.00135.

8. Ribas-Fernandes JJF, Solway A, Diuk C, McGuire JT, Barto AG, Niv Y, et al. A neural signature of hierarchical reinforcement learning. Neuron. 2011;71(2):370–379. doi: 10.1016/j.neuron.2011.05.042.

9. Yin HH, Ostlund SB, Knowlton BJ, Balleine BW. The role of the dorsomedial striatum in instrumental conditioning. Europearn Journal of Neuroscience. 2005;22(2):513–523. doi: 10.1111/j.1460-9568.2005.04218.x.

10. Daw ND, O’Doherty JP, Dayan P, Seymour B, Dolan RJ. Cortical substrates for exploratory decisions in humans. Nature. 2006;441(7095):876–879. doi: 10.1038/nature04766.

11. Mehlhorn K, Newell BR, Todd PM, Lee MD, Morgan K, Braithwaite VA, et al. Unpacking the exploration–exploitation tradeoff: A synthesis of human and animal literatures. Decision. 2015;2(3):191–215. doi: 10.1037/dec0000033.

12. Brovelli A, Laksiri N, Nazarian B, Meunier M, Boussaoud D. Understanding the Neural Computations of Arbitrary Visuomotor Learning through fMRI and Associative Learning Theory. Cerebral Cortex. 2008;18(7):1485–1495. doi: 10.1093/cercor/bhm198.

13. Jahanshahi M, Obeso I, Rothwell JC, Obeso JA. A fronto–striato–subthalamic–pallidal network for goal-directed and habitual inhibition. Nature Reviews Neuroscience. 2015;16(12):719–732. doi: 10.1038/nrn4038.

14. Caligiore D, Arbib MA, Miall CR, Baldassarre G. The super-learning hypothesis: Integrating learning processes across cortex, cerebellum and basal ganglia. Neuroscience and Biobehavioral Reviews. 2019;100:19–34. doi: 10.1016/j.neubiorev.2019.02.008.

15. Helmholtz H. Concerning the perceptions in general. In: Southall J, editor. Treatise on physiological optics (3rd ed., Vol. III, Translation 1962). New York: Dover; 1866. p. 214–230.

16. Dayan P, Hinton GE, Neal RM, Zemel RS. The helmholtz machine. Neural computation. 1995;7(5):889–904.

17. Doya K, Ishii S, Pouget A, Rao RPN, editors. The Bayesian Brain: Probabilistic Approaches to Neural Coding. Cambridge, MA: MIT Press; 2007.

18. Friston K. The free-energy principle: a unified brain theory? Nature Reviews Neuroscience. 2010;11(2):127–138. doi: 10.1038/nrn2787.

19. Griffiths TL, Kemp C, Tenenbaum JB. Bayesian models of cognition. Cambridge, UK: Cambridge University Press; 2008.

20. Toussaint M, Storkey A. Probabilistic inference for solving discrete and continuous state Markov Decision Processes. In: Proceedings of the 23rd international conference on Machine learning. ACM; 2006. p. 945–952.

21. Botvinick M, Toussaint M. Planning as inference. Trends in Cognitive Sciences. 2012;16(10):485–488. doi: 10.1016/j.tics.2012.08.006.

22. Kappen HJ, Gómez V, Opper M. Optimal control as a graphical model inference problem. Machine learning. 2012;87(2):159–182.

23. Bishop CM. Pattern recognition and machine learning. springer; 2006.

24. Kappel D, Nessler B, Maass W. STDP Installs in Winner-Take-All Circuits an Online Approximation to Hidden Markov Model Learning. PLoS Computational Biology. 2014;10(3):e1003511. doi: 10.1371/journal.pcbi.1003511.

25. Rao RP, Olshausen BA, Lewicki MS. Probabilistic models of the brain: Perception and neural function. Boston, MA: MIT press; 2002.

26. Jones M, Love BC. Bayesian Fundamentalism or Enlightenment? On the explanatory status and theoretical contributions of Bayesian models of cognition. Behavioral and Brain Sciences. 2011;34(4):169–88; disuccsion 188–231. doi: 10.1017/S0140525X10003134.

27. Maass W. Networks of spiking neurons: the third generation of neural network models. Neural networks. 1997;10(9):1659–1671.

28. Buesing L, Bill J, Nessler B, Maass W. Neural Dynamics as Sampling: A Model for Stochastic Computation in Recurrent Networks of Spiking Neurons. PLoS Computational Biology. 2011;7(11):e1002211. doi: 10.1371/journal.pcbi.1002211.

29. Orhan AE, Ma WJ. Efficient probabilistic inference in generic neural networks trained with non-probabilistic feedback. Nature communications. 2017;8:138. doi: 10.1038/s41467-017-00181-8.

30. Pouget A, Beck JM, Ma WJ, Latham PE. Probabilistic brains: knowns and unknowns. Nature Neuroscience. 2013;16(9):1170–1178. doi: 10.1038/nn.3495.

31. Maass W. On the computational power of winner-take-all. Neural computation. 2000;12(11):2519–2535.

32. Nessler B, Pfeiffer M, Buesing L, Maass W. Bayesian Computation Emerges in Generic Cortical Microcircuits through Spike-Timing-Dependent Plasticity. PLoS Computational Biology. 2013;9(4):e1003037. doi: 10.1371/journal.pcbi.1003037.

33. Bill J, Buesing L, Habenschuss S, Nessler B, Maass W, Legenstein R. Distributed Bayesian Computation and Self-Organized Learning in Sheets of Spiking Neurons with Local Lateral Inhibition. PLOS ONE. 2015;10(8):e0134356. doi: 10.1371/journal.pone.0134356.

34. Rückert EA, Neumann G, Toussaint M, Maass W. Learned graphical models for probabilistic planning provide a new class of movement primitives. Frontiers in Computational Neuroscience. 2013;6. doi: 10.3389/fncom.2012.00097.

35. Rueckert E, Kappel D, Tanneberg D, Pecevski D, Peters J. Recurrent Spiking Networks Solve Planning Tasks. Scientific Reports. 2016;6(1). doi: 10.1038/srep21142.

36. Tanneberg D, Paraschos A, Peters J, Rueckert E. Deep spiking networks for model-based planning in humanoids. In: Humanoid Robots (Humanoids), 2016 IEEE-RAS 16th International Conference on. IEEE; 2016. p. 656–661. Available from: http://ieeexplore.ieee.org/abstract/document/7803344/.

37. Daw ND, Niv Y, Dayan P. Uncertainty-based competition between prefrontal and dorsolateral striatal systems for behavioral control. Nature Neuroscience. 2005;8(12):1704–1711. doi: 10.1038/nn1560.

38. Viejo G, Khamassi M, Brovelli A, Girard B. Modeling choice and reaction time during arbitrary visuomotor learning through the coordination of adaptive working memory and reinforcement learning. Frontiers in Behavioral Neuroscience. 2015;9. doi: 10.3389/fnbeh.2015.00225.

39. Klein RM. Inhibition of return. Trends in Cognitive Sciences. 2000;4(4):138–147. doi: 10.1016/S1364-6613(00)01452-2.

40. Neal RM, Hinton GE. A view of the EM algorithm that justifies incremental, sparse, and other variants. In: Learning in graphical models. Springer; 1998. p. 355–368.

41. Bishop CM. Pattern recognition and machine learning. Berlin: Springer; 2006.

42. Jolivet R, Rauch A, Lüscher HR, Gerstner W. Predicting spike timing of neocortical pyramidal neurons by simple threshold models. Journal of computational neuroscience. 2006;21(1):35–49.

43. Dan Y, Poo Mm. Spike timing-dependent plasticity of neural circuits. Neuron. 2004;44(1):23–30.

44. Feldman D. The Spike-Timing Dependence of Plasticity. Neuron. 2012;75(4):556–571. doi: 10.1016/j.neuron.2012.08.001.

45. Markram H, Gerstner W, Sjöström PJ. Spike-Timing-Dependent Plasticity: A Comprehensive Overview. Frontiers in Synaptic Neuroscience. 2012;4. doi: 10.3389/fnsyn.2012.00002.

46. Zappacosta S, Mannella F, Mirolli M, Baldassarre G. General differential Hebbian learning: Capturing temporal relations between events in neural networks and the brain. Plos Computational Biology. 2018;14(8):e1006227. doi: doi.org/10.1371/journal.pcbi.1006227.

47. Kingma DP, Welling M. Auto-Encoding Variational Bayes; 2013.

48. Goodfellow I, Bengio Y, Courville A. Deep Learning. Boston, MA: The MIT Press; 2017. Available from: http://www.iro.umontreal.ca/~bengioy/dlbook.

49. Jaeger H. The ‘echo state’ approach to analysing and training recurrent neural networks-with an erratum note. Bonn, Germany: German National Research Center for Information Technology; 2001. 48.

50. Maass W, Natschläger T, Markram H. Real-time computing without stable states: a new framework for neural computation based on perturbations. Neural Comput. 2002;14(11):2531–2560. doi: 10.1162/089976602760407955.

51. Gibson JJ. The Ecological Approach to Visual Perception. Boston, MA: Houghton Mifflin; 1979.

52. Baldassarre G, Lord W, Granato G, Santucci VG. An embodied agent learning affordances with intrinsic motivations and solving extrinsic tasks with attention and one-step planning. Frontiers in Neurorobotics. publ2019;13(45). doi: 10.3389/fnbot.2019.00045.

53. O’Reilly RC, Frank MJ. Making working memory work: a computational model of learning in the prefrontal cortex and basal ganglia. Neural Computation. 2006;18(2):283–328. doi: 10.1162/089976606775093909.

54. Mannella F, Mirolli M, Baldassarre G. Goal-Directed Behavior and Instrumental Devaluation: A Neural System-Level Computational Model. Frontiers in Behavioral Neuroscience. 2016;10(181):e1–27. doi: 10.3389/fnbeh.2016.00181.

55. Brovelli A, Badier JM, Bonini F, Bartolomei F, Coulon O, Auzias G. Dynamic reconfiguration of visuomotor-related functional connectivity networks. Journal of Neuroscience. 2017;37(4):839–853.

56. Brovelli A, Chicharro D, Badier JM, Wang H, Jirsa V. Characterization of Cortical Networks and Corticocortical Functional Connectivity Mediating Arbitrary Visuomotor Mapping. Journal of Neuroscience. 2015;35(37):12643–12658. doi: 10.1523/JNEUROSCI.4892-14.2015.

57. Kriegeskorte N, Mur M, Bandettini PA. Representational similarity analysis – Connecting the branches of systems neuroscience. Frontiers in systems neuroscience. 2008;2:4.

